# Impact of Vitamin D on Gene Expression in Atlantic Salmon Skin and Potential Immunomodulation Against Salmon Louse Infection

**DOI:** 10.1101/2025.09.01.673453

**Authors:** Lewis D. Taylor, Courtney E. Gorman, Fintan Egan, Neil M. Ruane, Philip McGinnity, C. Darrin Hulsey

**Affiliations:** School of Biology and Environmental Science, University College Dublin, Belfield, Dublin, Ireland; Marine Institute, Newport, Co. Mayo, Ireland; School of Biological, Earth and Environmental Sciences, University College Cork, Cork, Ireland

**Keywords:** *Salmo salar*, *Lepeophtherius salmonis*, ectoparasite, immunomodulation, vitamin D

## Abstract

Vitamin D is a key micronutrient in vertebrate health that influences musculoskeletal function and likely modulates immune responses in tissues such as the skin. In aquaculture, Atlantic salmon (*Salmo salar*) are particularly vulnerable to skin pathogens, notably the ectoparasitic salmon louse (*Lepeophtheirus salmonis*). As the skin represents the first line of immunological defence, its capacity to mediate host-pathogen interactions may be influenced by dietary vitamin D. This study investigated the transcriptomic response of salmon skin following six months of dietary vitamin D supplementation, with a focus on immune-related gene expression. A total of 113 differentially expressed genes (DEGs) were identified, 46 of which are implicated in inflammation, immune signalling, and both innate and adaptive immunity. These DEGs were further compared to published RNA-seq data of salmon skin challenged with *L. salmonis*. Notably, while pro-inflammatory genes were regulated in opposing directions, heat shock proteins, lysozyme and antimicrobial peptides were consistently upregulated under both conditions. These findings suggest that vitamin D may modulate cutaneous immune responses by dampening inflammation and enhancing innate defences, potentially improving resistance to skin-associated pathogens such as salmon lice.

## 1. Introduction

The skin serves as both a defensive barrier and a vital interface between an organism and its environment. In humans, the skin forms the first line of immune defence against pathogens and facilitates the production of the essential nutrient vitamin D3 in response to solar UV radiation (White, 2013; Cashman, 2015; Dunlop *et al*., 2013). However, at northern latitudes, limited sunlight during winter restricts natural vitamin D synthesis, necessitating dietary intake from vitamin D-rich sources such as Atlantic salmon (*Salmo salar*) (Cui *et al*., 2022; Dimitrov *et al*., 2021). Atlantic salmon are among the richest natural sources of vitamin D (Moxness Reksten *et al*., 2022), and like humans, their vitamin D levels fluctuate in response to environmental factors (Jakobsen *et al., 2019;* Zerofsky *et al*., 2016). Unlike humans, Atlantic salmon obtain vitamin D exclusively from their diet (Cheng *et al*., 2019). This makes Atlantic salmon ideal candidates for biofortification through dietary supplementation in aquaculture (Jakobsen *et al*., 2019; Gorman *et al*., 2025).

Vitamin D plays a critical role in health and modulates immune responses in many vertebrates (Martens *et al*., 2020). Vitamin D deficiency is well-known for inducing long-term conditions like rickets, or the softening and weakening of bones, in a wide range of animals like birds, reptiles, ungulates, carnivores, and primates (Ismailova and White, 2022; Cashman *et al*., 2017; Uhl, 2018; Kumar, 2018; Chesney, 2010; Fiennes, 1974). However, in addition to its direct impacts on the skeletal system, vitamin D deficiency is linked to a wide range of short-term health conditions. In terrestrial vertebrates, vitamin D can mediate respiratory infections (Kumar, 2018; Walker and Modlin, 2009), inflammatory skin disorders (Zarei *et al*., 2019) and susceptibility to parasites like head lice (Khalid Awaad *et al*., 2023) and leishmaniasis (Martori *et al*., 2021). In some fish, like rainbow trout (*Oncorhynchus mykiss*), insufficient vitamin D is known to weaken skin integrity (Rider *et al*., 2023), increasing susceptibility to damage and infection (Zerofsky *et al*., 2016). Vitamin D could boost the immune responses of the skin of Atlantic salmon both in freshwater and saltwater environments.

Vitamin D’s immunomodulatory effects are largely driven by its regulation of immune-related gene expression (Ismailova and White, 2022; Martins *et al*., 2020). These genes fall into major functional categories, such as innate and adaptive immunity, inflammation, and immune signalling (Daryabor *et al*., 2023; Athanassiou *et al*., 2022). In mammals, vitamin D regulates genes involved in innate immune responses that provide immediate but broad-spectrum protection from pathogens (Athanassiou *et al*., 2022; Zhang and Zhang, 2023). Additionally, vitamin D influences adaptive immune responses, which are slower to activate but highly pathogen-specific (Cantorna and Arora, 2023; Hardiyanti *et al*., 2024; Cronkite and Strutt, 2018). For example, vitamin D enhances the expression of genes responsible for lysozyme production as well as anti-microbial peptides (AMPs) that are key factors in both innate immunity and antigen presentation (Hardiyanti *et al*., 2024; Soto-Dávila *et al*., 2020; Zhang *et al*., 2021). Additionally, vitamin D regulates inflammatory processes by preventing excessive or chronic immune activation (Fernandez *et al*., 2023; Giangreco *et al*., 2015) and facilitating immune cell migration to infection sites (Karkeni *et al*., 2019; Barnes and Amir, 2017). While these effects have been studied extensively in mammals, vitamin D’s influence on immune gene expression in fish, such as Atlantic salmon, remains poorly understood.

Farmed Atlantic salmon are highly susceptible to several skin pathogens, including the ectoparasitic salmon louse (*Lepeophtherius salmonis*). Vitamin D biofortification could, in addition to supporting human health, beneficially boost the Atlantic salmon’s immune response to several skin-associated pathogens. Because the skin serves as the primary immunological barrier between Atlantic salmon and economically important pathogens like salmon lice, vitamin D fortification of Atlantic salmon skin could alter immune gene expression and beneficially mediate salmon host-pathogen interactions.

Pathogens and parasites, such as the skin-feeding salmon louse, pose a major challenge to sustainable Atlantic salmon aquaculture (Stige *et al*., 2024; Kragesteen *et al*., 2021; Besnier *et al*., 2014). As one of the most prolific ectoparasites affecting Atlantic salmon populations (Brakstad *et al*., 2019; Cai *et al*., 2022), salmon lice cause significant skin damage, reduce growth rates, increase mortality, and leave fish vulnerable to secondary bacterial infections during the saltwater phase of the Atlantic salmon life cycle (Brakstad *et al*., 2019). These infections result in substantial economic losses, costing aquaculture operations millions of Euros annually (Cai *et al*., 2022; Fjørtoft *et al*., 2020; Larsen and Vormedal, 2021). A major reason for these challenges is that Atlantic salmon struggle to mount an effective immune response against the parasite (Misund, 2019; Vollset *et al*., 2023; Zhang *et al*., 2023; Carvalho *et al*., 2020), and anti-parasitic treatments are becoming increasingly ineffective due to resistance (Liu and Bjelland, 2014; Bjørn *et al*., 2020). In salmonids, thousands of genes are differentially expressed in the skin in response to salmon louse infections, yet how immune-related genes regulated by vitamin D intersect with these genes remains unknown (Øvergård *et al*., 2023; Carmona-Antoñanzas *et al*., 2016; Dalvin *et al*., 2020). Given vitamin D’s role in regulating immune gene expression in other vertebrates, biofortification may offer a novel strategy to prepare fish before being introduced into saltwater aquaculture and enhance Atlantic salmon immunity against these types of skin-feeding pathogens, providing economic and welfare benefits for aquaculture.

This study investigates how vitamin D biofortification affects immune gene expression in Atlantic salmon skin. First, we employed RNAseq to examine how vitamin D treatment influences gene expression patterns in Atlantic salmon skin, providing insights into its broader physiological effects. Secondly, we explored whether vitamin D modulates the expression of genes associated with immune function. Finally, we evaluated if vitamin D supplementation alters the expression of genes and pathways involved in the immune response to salmon louse infection, offering a novel perspective on the intersection of nutrition and pathogen resistance in Atlantic salmon aquaculture.

## 2. Materials and methods

### 2.1. Experimental Populations

Atlantic salmon were raised according to detailed methods described previously in Gorman et al. (2025). Briefly, two experimental populations were established, one from eggs obtained from a Norwegian commercially farmed strain, the other using eggs acquired from a captive-bred Irish strain used for experimental ocean release (sea ranching) programmes. At the commencement of the first feeding (April 2023), that period when alevins transition from utilising yolk sac reserves to exogenous food, the fry were fed a standard commercial feed for one week before the experimental groups were transitioned to experimental feeds. Bio authorisation was granted by The University College Dublin Animal Research Ethics Committee, which approved the use of salmon in this study: Research Ethics Reference Number AREC-23-01-Hulsey

### 2.2. Vitamin D Experimental Feeds

The two experimental aquafeeds, as described in Gorman *et al*. (2025), included additional vitamin D (cholecalciferol) in addition to the vitamin D included in a vitamin-mineral premix at one of two levels: low (0 µg/kg) and high (1000 µg/kg). Both feeds had a basal composition of 50% crude protein and 21% crude lipid, differing only in the concentration of vitamin D3 (cholecalciferol). All feeds included a standard salmon feed base consisting of fishmeal LT (50.0%), lysine (0.6%), methionine (1.2%), krill, squid, and CPSP90 in a 4:2:4 ratio (10.0%), wheat gluten (10.0%), soybean protein concentrate (3.6%), fish oil (10.0%), canola and soybean oil in a 1:1 ratio (2.5%), soybean lecithin (1.0%), choline chloride (0.5%) and betaine (0.5%). The vitamin-mineral premix (2.0%) contained (quantities per kg of premix): Sodium chloride, potassium chloride, calcium carbonate (support ingredients), butylated hydroxytoluene (20,000 mg), colloidal silica (176,700 mg), sepiolite (370,600 mg), D-calcium pantothenate (10,000 mg), biotin (300 mg), inositol (50,000 mg), anhydrous betaine (50,000 mg), iron sulphate monohydrate (600 mg), potassium iodide (50 mg), manganese (960 mg), zinc (750 mg), copper (900 mg), selenium (1 mg), vitamin B2 (3,000 mg), vitamin B12 (10 mg), niacinamide (20,000 mg), folic acid (1,500 mg), vitamin A (2,000,000 IU), vitamin E (10,000 mg), vitamin K3 (2,500 mg), vitamin B1 (3,000 mg), vitamin B6 / pyridoxine hydrochloride (2,000 mg), and vitamin D3 (200,000 IU),

All ingredients were mixed in a 10 L mixer, ground with a hammer mill (UPZ 100, Hosokawa-Alpine, Augsburg, Germany) to 0.5□mm. The diets were extruded using a five-section twin-screw extruder (Evolum 25, Clextral, Firminy, France), fitted with 0.5 mm and 2 mm die holes. The pellets were dried at 27 °C in a drying chamber (Airfrio, Almería) and cooled at room temperature. Experimental Diet Service at the University of Almeria (Spain) formulated the experimental feeds. The two experimental feed treatments were provided *ad libitum* to the respective tanks containing the developing alevins.

### 2.3. Sampling

The skin of the Atlantic salmon was sampled the week beginning October 1, 2023, after the fish had been fed experimental diets for approximately six months and were sacrificed with an overdose of MS222. Samples of the skin and the fillet were dissected from the dorsal right side of the fish. To sample the skin, a rectangular area of approximately 3 x 1 cm was removed, and the skin was then separated from the underlying muscle using a scalpel. The dissected skin samples were stored in RNAlater (Sigma-Aldrich) in 1.5 ml Eppendorf tubes before being sent for RNA sequencing. At the end of the experiment, the tissue concentration of vitamin D in the axial musculature (filet) taken directly underneath our skin samples was quantified through triple/quadrupole mass spectrometry and electrospray ionisation at the National Food Institute, Technical University of Denmark, as reported in Gorman *et al*. (2025).

### 2.4. Atlantic salmon Skin mRNA Sequencing

To generate RNAseq libraries for the skin samples, 12 samples were sent via dry ice to Novogene at their Cambridge Sequencing Centre (Cambridge, UK). Novogene conducted RNA extraction, mRNA isolation, RNAseq library preparation and sequencing. The initial raw data produced (fastq) was processed by Novogene using their in-house pipeline (Perl scripts). Clean reads were acquired by removing any reads that had poly-Ns and showed signs of low quality. All downstream analyses used high-quality reads. The Atlantic salmon (Ssal_v3.1) was used as the reference genome and annotation model.

### 2.5. Differential Expression Quantification – Vitamin D (low versus high)

The cleaned fastq files produced from Novogene were run through the Nextflow rnaseq pipeline (23.04.3) using nf-core (3.11.2) (Patel *et al*., 2024). The resulting count tables were employed for differential expression analysis using the DESeq2 R package 1.20.0 (Love *et al*., 2014). For this experiment, significance was determined using Benjamini-Hochberg adjusted p-values (p-adjusted) rather than raw p-values and a log fold change of ≤ −1 or ≥ 1. These adjusted p-values account for the high number of statistical comparisons made on the same treatments when performing differential expression analysis, which can lead to increased false discovery rates (Benjamini and Hochberg, 1995). The data set was prefiltered to exclude loci with minimum counts < 10 and that were present in less than five samples to reduce the number of low and zero-count genes. Genes were determined to be differentially expressed if the p-adjusted values were less than or equal to 0.05. The DESeq2 design included vitamin D treatment as the condition while controlling for strain as a batch effect. Differentially expressed genes identified from vitamin D treatment groups were annotated from Ensembl identifiers using the biomaRt package from the ssalar_gene_ensembl database (Durinck *et al*., 2009).

### 2.6. Differential Expression Quantification – Sea Louse (uninfected versus infected)

Previously published data from Øvergård *et al*. (2023) were used to compare differentially expressed genes in salmon louse-infected Atlantic salmon skin with Atlantic salmon skin from our vitamin D treatments. Because we thought it was most biologically relevant to compare to our vitamin D treatments, we only included the contrast between uninfected Atlantic salmon (n = 5) and Atlantic salmon infected with salmon louse (n = 5) skin samples taken directly under the sight of sea louse feeding sites (Øvergård *et al*., 2023). The salmon louse data, which was previously evaluated using a different genome, was reanalysed here using the Atlantic salmon genome Ssal3.1 to facilitate comparisons. As for the vitamin D data, the salmon louse infection data were run through Nextflow’s rnaseq pipeline, but the fastq files were acquired from the Sequence Read Archive NCBI database (Sequence Read Archive, 2009). The files were converted into count tables using the tximport version 3.19 (Soneson *et al*., 2016). These count files were then analysed with DESeq2 (Love *et al*., 2014). To ensure parity of the comparisons, the salmon louse data were prefiltered using the same criteria we used with our data.

### 2.7. Identification of immune genes

Differentially expressed genes from experiments were annotated in several ways to determine if they likely had an immune function. The genes from each group were first annotated using the AnnotationDbi (version 3.19) package from (Soneson *et al*., 2016), Bioconductor version 1.66.0 (Pagès *et al*., 2024). Genes without annotated Ensembl IDs were subjected to a BLAST (Marini *et al*., 2020) search of the longest exon to determine if there was a match on NCBI. Any genes that were not initially named on Ensembl but found substantial matches or homologues on BLAST were annotated to further determine their putative function. The matching percentage was set at a minimum of 80% and coverage of 80%. Genes that had names and possessed an accession number were then searched for using the Web of Science with search terms gene symbol AND Immune*. The complete list of annotated genes is provided (Supplementary Table 1), but the genes associated with immunity are primarily the focus of discussion below. The immune gene categories were determined using Web of Science (Web of Science™) using the individual gene names and the key phrases ‘innate immune system’, ‘adaptive immune system’, ‘inflammation’, and ‘immune signalling’.

### 2.8. Gene Set Enrichment Analysis

Gene set enrichment analysis (GSEA) of our vitamin D skin and the salmon louse data was conducted using the clusterProfiler package in R (Yu, 2024; Altschul *et al*., 1990) to identify putative functions and underlying biological phenomena associated with all expressed genes identified from DESeq2 (Altschul *et al*., 1990; Subramanian *et al*., 2005). GSEA was examined using the full DESeq2 output as it aggregates the per-gene statistics across genes within a gene set, thereby making it possible to detect situations where expression changes in a small but coordinated way. The annotation used for acquiring the GO terms was AH114250, the most current annotation for Atlantic salmon at the time of analysis. For the GSEA, significance was determined by an adjusted p-value of ≤ 0.05.

### 2.9. Statistical Analysis

Vitamin D concentration for low and high treatment groups as determined by triple/quadrupole mass spectrometry was analysed using a one-way ANOVA, with p□≤□0.05 considered significant and □p ≤□0.001 being considered highly significant.

## 3. Results

### 3.1. Vitamin D tissue concentration

There was a significant (p□≤□0.001) difference in vitamin D concentration assimilated in the Atlantic salmon filets between the two treatments, with the low group average of 860 ng/mg in the low treatment and 1840 ng/mg in the high treatment groups.

### 3.2. RNA sequencing libraries

Once cleaned, the RNAseq libraries from the high (n = 6) and low (n = 6) vitamin D groups had a mean size of 13.1 Gb and ranged from 10.38 to 16.17 Gb (Supplementary Table 2). The mean total mapping rate of the libraries was 86.46% and ranged from 71.35% to 91.51% (Supplementary Table 2). The SRA GenBank accession numbers ranged as follows: SRA41355584-SRA41355595.

### 3.3. Vitamin D-induced differentially expressed genes

Of the total 33195 genes inferred to be expressed, 113 genes were found to be significantly differently expressed (adjusted p-value ≤ 0.05) between the low and high vitamin D treatment groups (Fig. 2) in the skin. Of the significant genes, 101 were downregulated, and 12 were upregulated. Of these, 23 novel genes could not be annotated (Supplementary Table 3), which left 90 genes that we examined more fully. These included a number of genes not clearly related to immunity, such as *mrps9* (mitochondrial ribosomal protein S9)*, nadk2* (NAD kinase 2) and *tars2* (threonyl-tRNA synthetase 2) that are all most commonly associated with mitochondrial function. Several other genes have diverse functions such as *krt18a.1* (keratin 18a) that could contribute to scale formation and repair, and *calml4* (calmodulin like 4) that is likely involved in calcium signalling. Based on the levels of gene expression as measured by log_2_ fold change for all genes in the vitamin D manipulated Atlantic salmon skin (Supplementary Table 1), the genes with the greatest up- and downregulation were *igkv* (Immunoglobulin kappa chain V region) (log_2_ fold change 4.46) and *esf1* (ESF1 nucleolar pre-rRNA processing protein homolog) (log_2_ fold change −5.46).

**Fig. 1.**
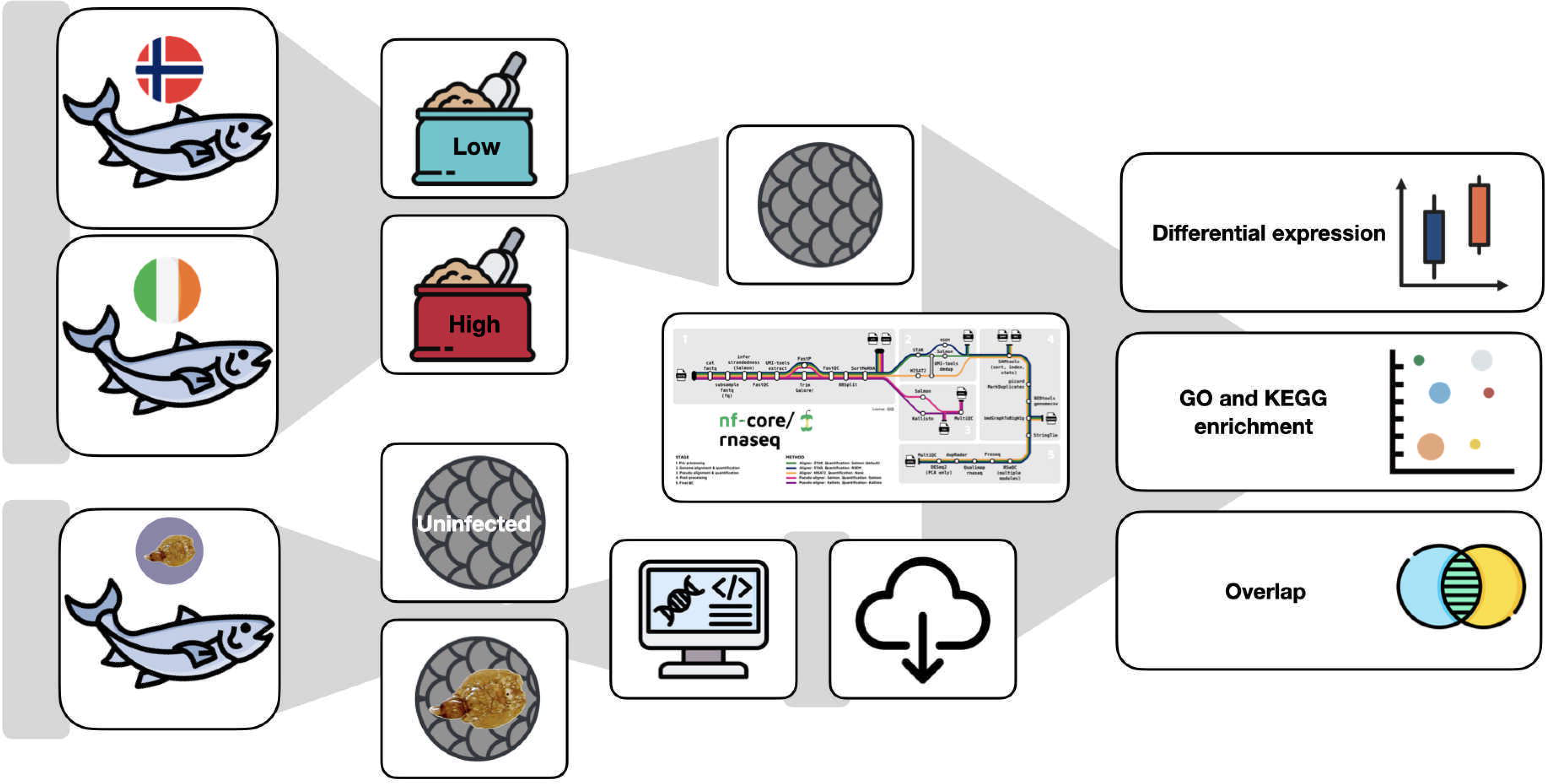
Experimental Workflow for Skin RNA-seq Analysis of our Vitamin D Supplementation and Published Salmon Louse Infection in Atlantic Salmon. The Atlantic salmon RNAseq data were derived from two separate experiments: 1. vitamin D supplementation and 2. salmon uninfected or exposed to salmon louse infection (41). Vitamin D treatment groups received either low or high doses and then the skin was removed and RNA sequenced. Both sets of data were analysed using the same pipeline.

**Fig. 2.**
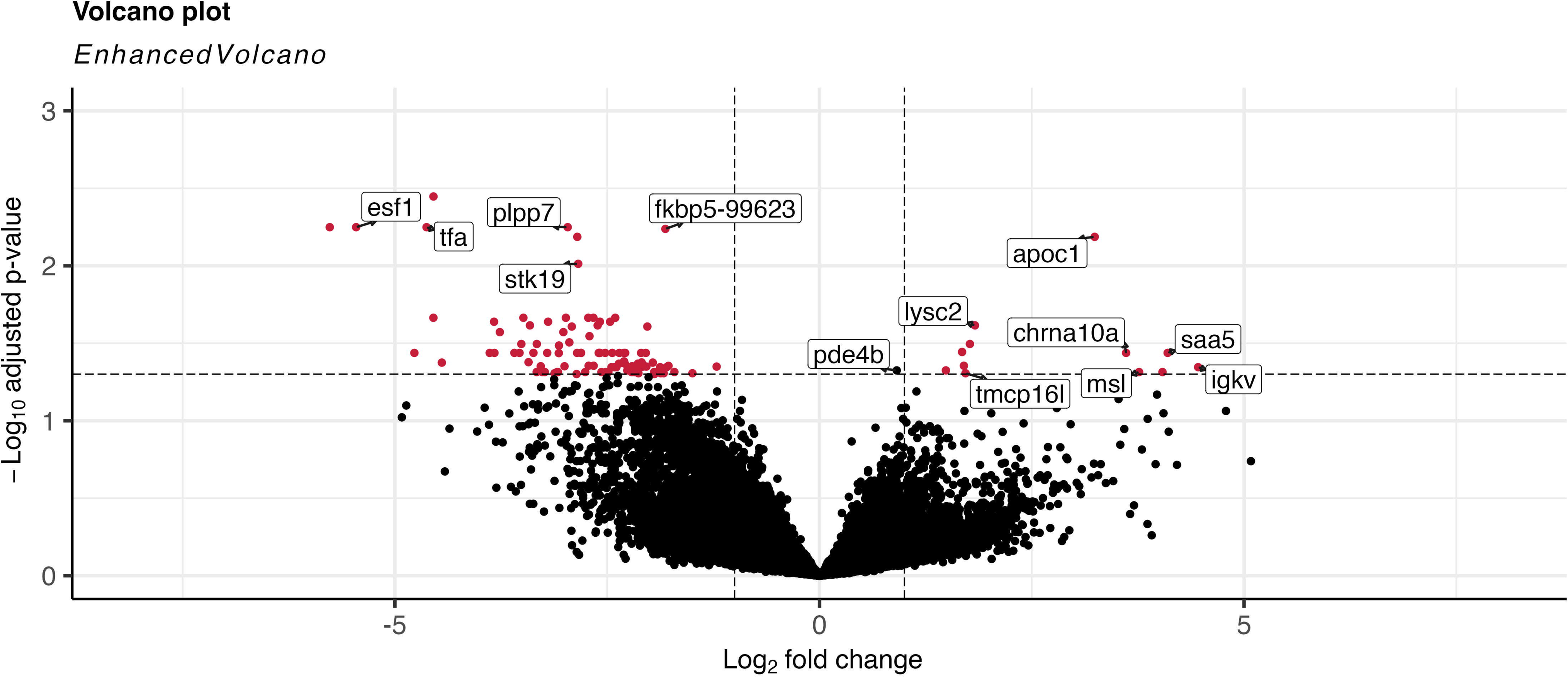
Volcano plot of differentially expressed genes between the high-treatment and low-treatment groups. The red data points are genes with both a significant (adjusted p-value ≤ 0.05) and log_2_ fold change value of ≥ 1 or ≤ −1. These genes represent the difference in expression between high and low vitamin D treatment groups. The black data points are genes that did not meet these thresholds. The genes with log_2_ fold change ≥ 1 are upregulated, and genes with ≤ −1 are considered downregulated. The labelled dots represent annotated genes with the greatest identifiable expression.

### 3.4. Immune Gene Expression

We identified 45 putative immune genes from the differential expression analysis and enhanced these annotations by analysing the sequences using BLAST to identify novel genes and their putative functions (Supplementary Table 4). To facilitate comparisons and discussion, our vitamin D-modulated expression of immune-related genes were grouped into four key functional categories: innate immunity, adaptive immunity, inflammation and immune signalling (Fig. 3). Concerning innate immunity, genes such as *bpifcl* (bactericidal/permeability-increasing fold containing family C, like), *cats* (cathepsin S) and *lysc2* (lysozyme), which are involved in antimicrobial defence and lysozyme activity, were upregulated. Within adaptive immunity, we observed an upregulation of *igkv* and a downregulation of *igh-a* (immunoglobulin heavy chain locus A), both associated with immunoglobulins’ structure and function in Atlantic salmon. In the inflammation category, vitamin D supplementation was associated with the downregulation of several pro-inflammatory genes such as *fkbp5-99623* (FKBP prolyl isomerase 5)*, fkbp5-79897* and *ptgs2a* (prostaglandin-endoperoxide synthase 2a) (Supplemental Table 4). Additionally, there was downregulation of *tfa* (transferrin a), which functions as a Fe²□ transporter, and upregulation of anti-inflammatory genes such as *chrna10a* (cholinergic receptor, nicotinic, alpha 10a) and *pde4b* (phosphodiesterase 4B). For immune signalling, altered expression of genes like *foxj2* (forkhead box J2) and *arhgap12* (rho GTPase-activating protein 12) highlighted potential roles in regulating the migration of immune cells to sites of infection or injury. Four genes, *tsc22d4* (TSC22 domain family protein 4-like)*, rbbp4* (histone-binding protein RBBP4-like)*, slc43a3a* (solute carrier family 43 member 3a), and *ankrd6* (ankyrin repeat domain-containing protein 6), were identified as being involved in modulating immune cell signalling. Additional genes that are likely involved in immune function but did not fit well into the above categories were also differentially expressed. For instance, the antiparasitic gene *apoc1* (Apolipoprotein C-I) and the *stk19* (serine/threonine-protein kinase 19) gene are associated with MHC (major histocompatibility complex) function. Finally, genes such as *vlgtp1* (interferon-induced very large GTPase 1-like)*, arih1* (E3 ubiquitin-protein ligase arih1-like), and *serpinc1* (serpin peptidase inhibitor, clade C) play roles in interferon stimulation and were altered by the vitamin D treatments.

**Fig. 3.**
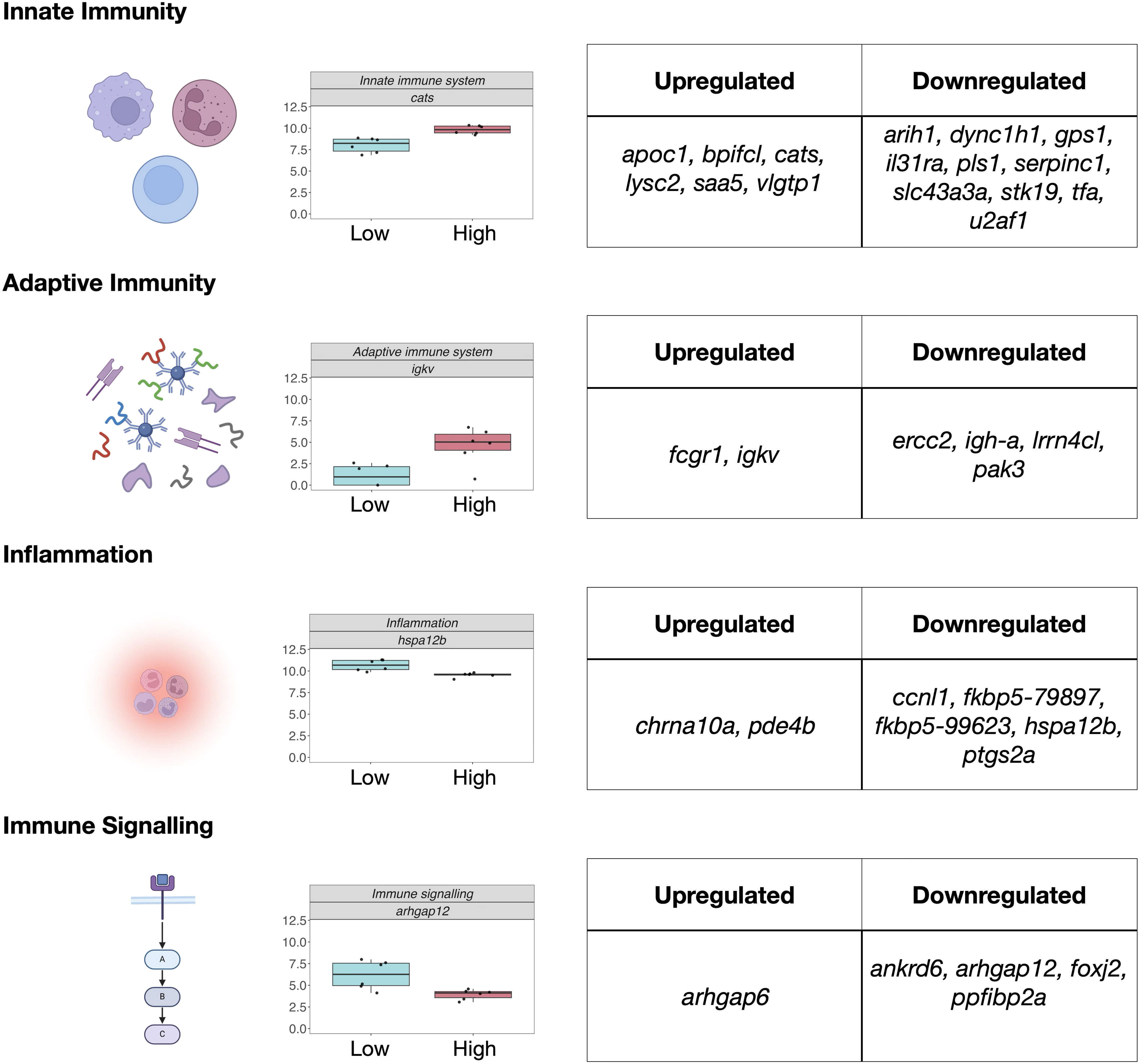
Expression of immune-related genes in Atlantic salmon skin following vitamin D supplementation. Box plots display the normalised log□ expression levels of representative immune-related genes in the skin of Atlantic salmon treated with low (cyan) and high (red) doses of vitamin D. Based on the literature, the genes are grouped into four major functional categories: innate immunity upregulated: *apoc1, bpifcl, cats, lysc2, saa5, vlgtp1*; downregulated: *arih1, dync1h1, gps1, il37ra, pls1, serpinc1, slc43a3a, stk19, tfa, u2af1*. Adaptive immunity *upregulated*: *fcgr1, igkv; downregulated: ercc2, igh-a, lrrn4cl, pak3*. Inflammation upregulated: *chrna10a, pde4b*; downregulated*: ccnl1, fkbp5-79897, fkbp5-99623, hspa12b, ptgs2a*. Immune signalling upregulated: arhgap6; downregulated*: ankrd6, arhgap12, foxj2* and *ppfibp2a*. Other notable immune related genes that did not fit clearly into the four categories were *ubtd1, ankrd9, rbbp4, tcb1, ccdc83, tsc22d4*, and *mcm10*.

### 3.5. Gene Expression Overlap Between Salmon Louse Infection and Vitamin D

We found several genes differentially expressed in the skin in response to vitamin D that could intersect with salmon louse infection (Fig. 4). In response to salmon louse infection, 2,617 genes were significantly differentially expressed (adjusted p ≤ 0.05). By contrast, only 90 annotated genes were differentially expressed due to vitamin D treatment, with an overlap of seven differentially expressed genes in both the salmon louse-infected and vitamin D-fortified Atlantic salmon skin.

**Fig. 4.**
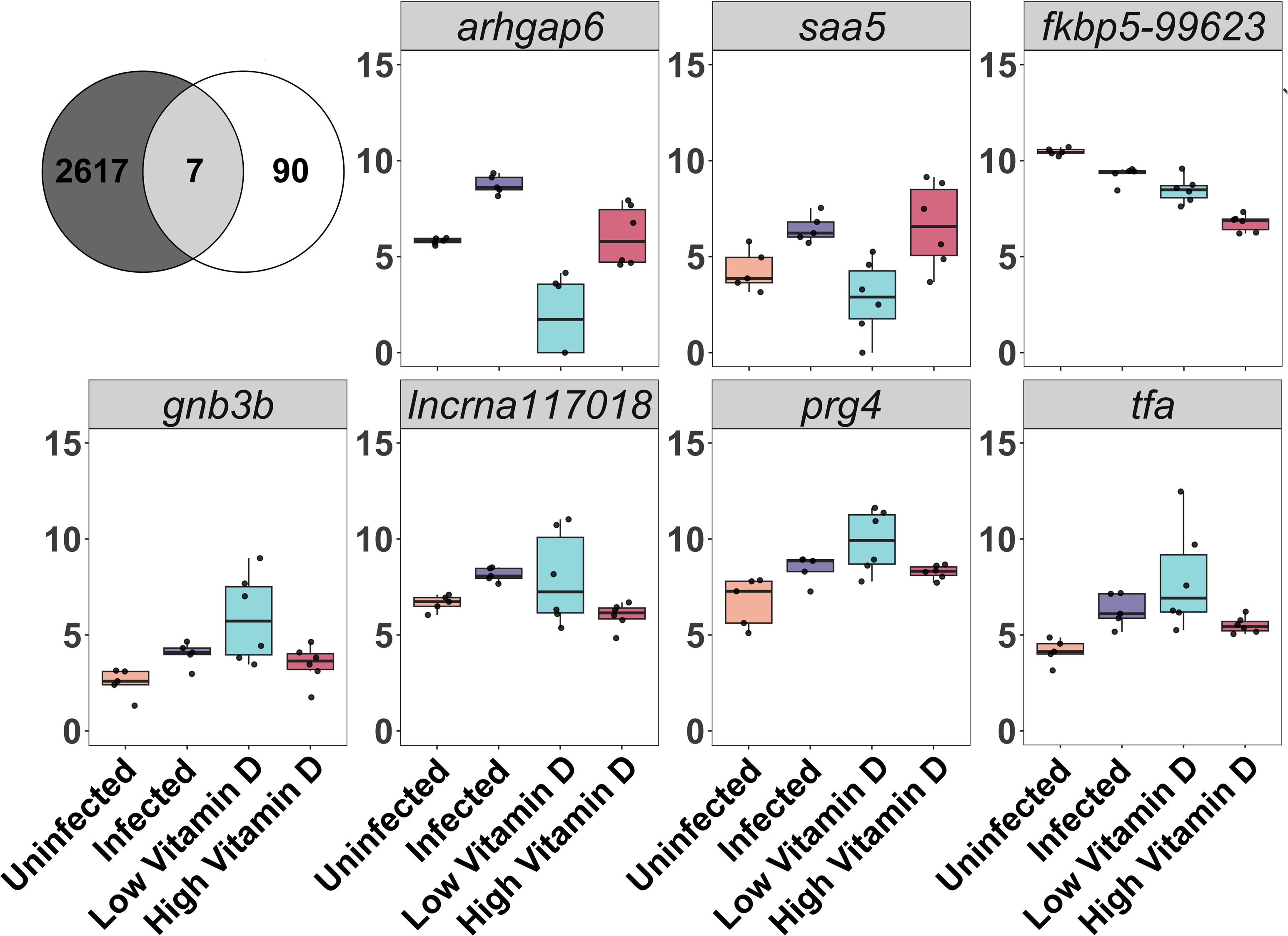
Differential gene expression in Atlantic salmon in response to salmon louse infection and Vitamin D treatment. (A). Venn diagram showing intersections of differentially expressed genes between salmon louse infection and vitamin D treatment. A total of 2617 genes were significantly differentially expressed in infected fish (adjusted p-value ≤ 0.05), with 90 of these annotated genes modulated by vitamin D treatment. Genes were filtered based on a log2 fold change threshold (≤ −1 or ≥ 1).(B) Box plots of normalised gene expression (log2-transformed) for selected genes *gnb3b*, *lncrna117018*, *prg4*, *tfa*, *arhgap6*, *saa5* and *fkbp5-99623* across experimental conditions: uninfected (orange), infected (purple), low-dose vitamin D treatment (cyan) and high-dose vitamin D treatment (red).

The seven genes that overlapped were *arhgap6* (Rho GTPase activating protein 6), *saa5* (Serum amyloid A-5 protein)*, fkpb5-99623, gnb3b* (guanine nucleotide binding protein (G protein), beta polypeptide 3b)*, lncrna117018* (lncRNA)*, prg4* (Proteoglycan 4) and *tfa*. The overlapping genes between the vitamin D-fortified groups and the salmon louse-infected groups exhibited various patterns in their direction of gene expression. Notably, *arhgap6* expression was upregulated in infected Atlantic salmon compared to uninfected controls and was upregulated in the high vitamin D treatment group compared to the low treatment groups but still lower than the salmon louse groups. A similar pattern of expression can be seen with *saa5*, where the salmon louse-infected and the high vitamin D group were both upregulated compared to the uninfected and low vitamin D groups. Conversely, *fkbp5-99623* expression was downregulated in the high vitamin D group and was downregulated in the infected group compared to the uninfected Atlantic salmon. Distinct expression patterns were also observed for *gnb3b*, which exhibited upregulation in the infected Atlantic salmon and downregulation in the high vitamin D treatment group. Similarly, the long non-coding RNA *lncrna117018* showed upregulation in the infected Atlantic salmon and downregulation in the higher dose of vitamin D. The expression of *prg4* was upregulated in the infected Atlantic salmon and downregulated in the high vitamin D group. Finally, *tfa* was upregulated in the salmon louse-infected Atlantic salmon and was downregulated in the high vitamin D group.

### 3.6. Gene Set Enrichment Analysis of Vitamin D Treatment Groups

We found 59 GO categories that were enriched in our GSEA of the Atlantic salmon skin in response to vitamin D. For the vitamin D treated Atlantic salmon, the enrichment analysis found there were 15 matches for biological processes, 12 for molecular function, and 6 for cellular components (Fig. 5A). The GO category with the greatest gene count was transmembrane signalling receptor activity. Ribosome activity was found to have the greatest gene ratio (0.6). Furthermore, GO terms associated with lysing activity and cytokine receptor activity exhibited notable overlaps. Several categories associated with the immune system were enriched (Supplementary Table 5).

**Fig. 5.**
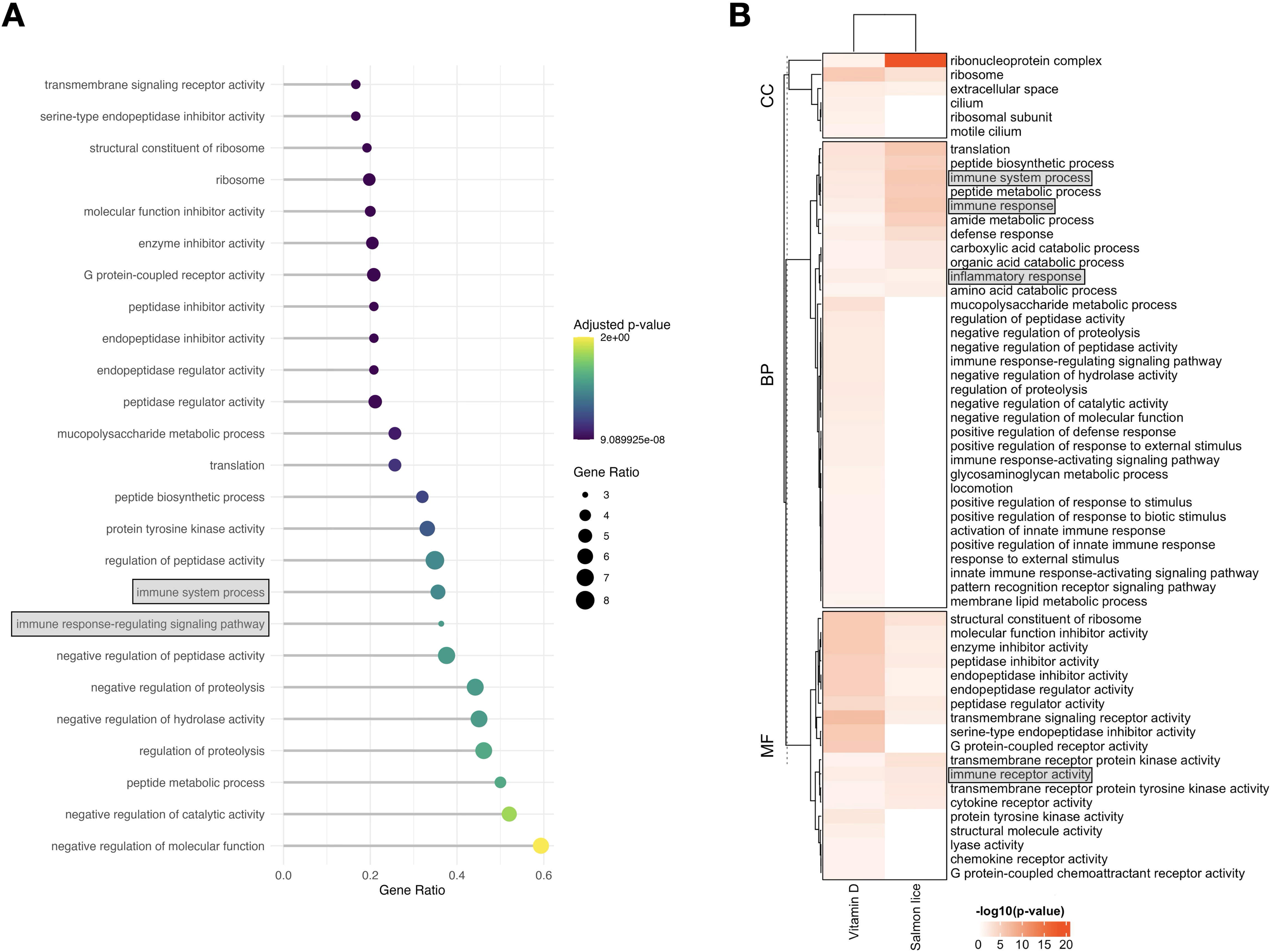
Gene Set Enrichment Analysis (GSEA) of significant genes in the skin between high vs low vitamin D treatment groups. (A). A dot plot displays the top 20 significantly enriched Gene Ontology (GO) terms for down regulated genes, including terms related to enzyme inhibitor activity, negative regulation of peptidase activity, and immune system processes. The size of the dots represents the gene ratio, while the colour gradient reflects the adjusted p-value, with shades ranging from green to purple, indicating increasing significance. (B). A hierarchically clustered heatmap showing enriched GO terms categorised into cellular component, biological process, and molecular function. The heatmap compares vitamin D modulation with salmon louse infection, highlighting pathways such as ribosome-related activity, immune system processes, and peptide metabolic processes. The shading intensity corresponds to -log10(adjusted p-value), with darker shades indicating higher significance.

### 3.7. Overlap between Vitamin D-fortified Atlantic salmon and Salmon louse-infected Atlantic salmon

The enriched GO categories in our skin data showed some overlap with those observed during salmon louse infection (Fig. 5). Specifically, the GSEA analysis of skin infected with Atlantic salmon lice revealed 27 overlapping biological processes, 14 overlapping molecular function terms, and six overlapping cellular component terms between the vitamin D treatments (Supplementary Table 6). Additionally, four prominent immune-related GO terms were shared between the vitamin D and salmon louse datasets: immune system process, immune response, inflammatory response, and immune receptor activity.

## 4. Discussion

Vitamin D was found here to impact gene expression in the skin of Atlantic salmon. These alterations could have functional consequences for immunity and interactions with pathogens. Vitamin D manipulation resulted in 113 differentially expressed genes in the skin (Fig. 2), several of which were involved in metabolism, cellular communication, inflammatory pathways, and the immune system. Vitamin D supplementation altered the expression of immune-related genes, including those involved in innate and adaptive immunity, inflammation, and immune signalling. Vitamin D also influenced the expression of several genes in Atlantic salmon skin previously associated with Atlantic salmon immunity, such as immunoglobulins and several anti-microbial peptides (AMPs) (Supplemental Table 4). In our evaluation of whether our vitamin D treatments affect genes whose expression is altered during salmon louse infections in Atlantic salmon, we identified seven overlapping differentially expressed genes. We also identified several gene pathways that overlapped and represented enriched immune categories. Vitamin D modulates immune gene expression in Atlantic salmon skin and may play a role in mediating many host-pathogen interactions.

Several genes that were differentially expressed in the skin in response to increased vitamin D have broad physiological effects (Fig. 5a). For instance, vitamin D influenced several mitochondrial-associated genes in Atlantic salmon skin, such as *mrps9, nadk2* and *tars2* (threonyl-trna synthetase 2, mitochondrial transcript) (Mootha *et al*., 2003; Jiang *et al*., 2024; Zhang and Zhang, 2023; Zheng *et al*., 2022). Vitamin D is known to regulate mitochondrial gene expression in Atlantic salmon muscle (Gorman *et al*., 2025), and finding these changes in the skin indicates that the effects of vitamin D on Atlantic salmon mitochondrial function may be more systemic. We also found that vitamin D influenced genes such as *krt18a.1*, which is important for epithelial tissues like skin and the linings of organs and blood vessels, and *calml4*, which is involved in calcium signalling (Ho *et al*., 2022; Lessard *et al*., 2015). These results suggest that the broad physiological effects and impact of vitamin D on general health, as observed in terrestrial vertebrates, likely extend to fish like Atlantic salmon.

Our vitamin D treatments impacted the expression of many genes in the skin that could broadly impact immune functions such as innate immunity, adaptive immunity, inflammation, and immune signalling (Athanassiou *et al*., 2022). Of the differentially expressed genes that we detected in response to vitamin D enhancement, almost half (45 genes) are known to be associated with various aspects of immunity that synergistically could help Atlantic salmon fight pathogens (Supplementary Table 4).

Of the differentially expressed immune genes identified, at least 16 have a prominent function in innate immunity, which acts as the first line of defence by responding rapidly to antigen-independent foreign bodies (Lessard *et al*., 2015). For instance, we found changes in the expression of three genes commonly associated with antimicrobial peptides (AMPs): *saa5, bpifcl, and apoc1* (Fig. 3). The *saa5* gene recruits immune cells to sites of inflammation, aids in identifying pathogens during infections (Tang *et al*., 2018), and can prevent secondary bacterial infections at sites of tissue trauma (Chi *et al*., 2024). The *bpifcl* gene is an AMP that specifically targets gram-negative bacteria, while *apoc1* is known for its bactericidal and defensive role against protozoan ectoparasites (Buks *et al*., 2023). Notably, we did not observe differential expression of cathelicidin, an AMP commonly associated with parasitic defence in Atlantic salmon (Øvergård *et al*., 2023). Yet, vitamin D changes in the expression of antimicrobial peptides that impact innate immunity could be particularly important for how Atlantic salmon respond to skin-associated pathogens.

Vitamin D likely influences Atlantic salmon’s innate and adaptive immune function through the differential expression of several immune-related genes in the skin. The upregulation of genes encoding lysozyme (*lysc2*) and cathepsin S (*cats*) highlights vitamin D’s role in bolstering innate immunity (Fig. 3). Lysozyme activity, which can be enhanced by *lysc2* expression, improves defences against bacteria (Zhang and Zhang, 2023). Increased lysozyme activity in response to vitamin D supplementation has also been observed in other teleost fishes (Zhang and Zhang, 2023). Together, these findings suggest that increased vitamin D levels could prime the immune system to favour innate immune responses and thereby potentially enhance the skin’s first-line defences against many pathogens that infect Atlantic salmon.

Several upregulated genes likely also influence adaptive immune components such as B cells, T cells and antigen-presenting cells (Rojas *et al*., 2024). For instance, we observed the upregulation of *igkv* and *fcgr1* (high-affinity immunoglobulin gamma Fc receptor I) alongside the downregulation of *igh-a* (Fig. 3). These genes are all integral to the structure and function of antibodies (McCormack *et al*., 2021; Bonilla and Oettgen, 2010; Johansson *et al*., 2023; Aslam *et al*., 2020). These results underscore the complex interactions between vitamin D and the adaptive immune response, which warrant further investigation to elucidate their implications for host defence in salmonids.

Several differentially expressed genes in the vitamin D treatments could also link innate and adaptive immunity to inflammation. For instance, vitamin D supplementation was associated with the downregulation of several pro-inflammatory genes, such as *fkbp5-99623, fkbp5-79897*, and *ptgs2a* (Fig. 5 and Supplementary Table 4). The two duplicates of the *fkbp5* gene are found on different chromosomes, but both likely function generally as immunophilins and play an important role in innate immune regulation (Zhang *et al*., 2016). The *fkbp5* gene is critical for the attachment and internalisation of viruses such as infectious Atlantic salmon anaemia virus (ISAV) in Atlantic salmon cells (Yasuike *et al*., 2010).

Reduced FKBP*5* activity lowers ISAV viral loads (Zhao *et al*., 2019). The reduced *ptgs2a* activity, which encodes prostaglandin-endoperoxide synthase 2, is known to occur with increased vitamin D levels (Giangreco *et al*., 2015; Gervais *et al*., 2022; Houston *et al*., 2020), and in salmonids, *ptgs2a* is an indicator of stress and disease (Liu *et al*., 2021). Additionally, anti-inflammatory genes such as *chrna10a* and *pde4b* were upregulated (Wang *et al*., 2014; Ji *et al*., 2019; Cashman *et al*., 2017). Excessive inflammation could impair growth and health in farmed Atlantic salmon, and our results support vitamin D’s established role in promoting an anti-inflammatory profile that could be conserved across vertebrates.

Our analysis of genes related to immune signalling (Fig. 4) also revealed altered expression in genes such as *foxj2* and *arhgap12*. The *foxj2* gene is part of the Forkhead box family, which is involved in tumorigenesis and suppresses the movement of immune cells into tumours (Scalavino *et al*., 2024; Richter *et al*., 2022). Similarly, the Rho GTPase-activating protein family member *arhgap12* plays a role in immune signalling and migration (Qiu *et al*., 2015; Fei *et al*., 2024). Vitamin D supplementation could readily influence immune cell trafficking and localisation to Atlantic salmon skin.

The overlap of particular differentially expressed genes found between vitamin D-fortified Atlantic salmon and those infected with the salmon louse (Fig. 4) suggests that vitamin D could indirectly affect gene expression related to salmon louse infections. For instance, there were clear changes in the gene *saa5* in the high-vitamin D group when compared with salmon louse-infected Atlantic salmon. Additionally, we identified differential expression of *lncrna117018* that encodes a novel long non-coding RNA in both datasets, and so this non-protein coding RNA should be further investigated for its role in Atlantic salmon immune reactions. The upregulation of the gene *prg4* (proteoglycan 4-like), in salmon louse-infected samples, suggests increased tissue repair activity (Fei *et al*., 2024), and its downregulation in high vitamin D-treated Atlantic salmon could help reduce sea louse-associated stress responses in the skin (Nanda *et al*., 2023). The downregulation of other inflammatory genes, such as *fkbp5-99623* and *ptgs2a*, further supports the hypothesis that vitamin D supplementation reduces inflammation while bolstering immune responses against bacterial infections. Despite directly affecting only a few genes associated with sea louse infection, vitamin D might have an outsized effect not only on host interactions with Atlantic salmon lice but could also likely improve survival rates in Atlantic salmon by reducing secondary bacterial and viral skin infections (Gervais *et al*., 2022; Scalavino *et al*., 2024).

There also appeared to be an overlap in immune pathways influenced by vitamin D treatment and those activated during salmon louse infections based on our GSEA analysis (Fig. 5B). Four immune-related GO terms were shared between the vitamin D and salmon louse datasets including immune system process, immune response, inflammatory response, and immune receptor activity. Notably, Atlantic salmon infected with the louse displayed increased expression of cellular components associated with immune system responses and processes that can be driven by infection (Larsen and Vormedal, 2021; Vollset *et al*., 2023). Our vitamin D-treated Atlantic salmon skin exhibited increased immune activity, and there was a clear overlap in responses, such as inflammatory activity, between the vitamin D treatment groups and salmon louse-infected fish. These findings underscore the potential for vitamin D to modulate both the Atlantic salmon immune response in the skin and the interaction between Atlantic salmon and pathogens in aquaculture.

Atlantic salmon biofortification with vitamin D in aquaculture could have multiple beneficial outcomes. Vitamin D can readily enhance the vitamin D content in fish farmed for human consumption (Jakobsen *et al*., 2019; Zerofsky *et al*., 2016; Gorman *et al*., 2025). Additionally, vitamin D augmentation could help the immune responses of Atlantic salmon to a number of pathogens. Our experiments suggest that fortifying Atlantic salmon in freshwater hatcheries prior to transfer to aquaculture facilities at sea could provide an immune boost to the fish. Priming several facets of their immune system with vitamin D could defend against several viruses, bacteria, and skin-mediated parasites like sea lice (Sadarangani *et al*., 2015; Salamony *et al*., 2025). Future studies should extend our analyses here and experimentally examine if the response of Atlantic salmon skin to vitamin D is similar in freshwater alevins and saltwater post-smolts. Additionally, it would be interesting to know if vitamin D effectively primes the skin of Atlantic salmon and allows them to resist commercially impactful parasites like sea lice more extensively. The addition of greater amounts of vitamin D to Atlantic salmon feed could be a win/win for both human consumption and the general immune health of Atlantic salmon in aquaculture.

## Acknowledgements

This study was funded by a Science Foundation Ireland Frontiers for the Future grant (21/FFP-P/10171). PMcG was supported by the Science Foundation Ireland Investigators Programme (SFI/15/IA/3028), by Science Foundation Ireland in conjunction with the Biotechnology and Biological Science Research Council (UK) grant award (16/BBSRC/3316) and by the Marine Institute (RESPI/FS/20/01).

