## Supplementary material for "Impact of Vitamin D on Gene Expression in Atlantic Salmon Skin and Potential Immunomodulation Against Salmon Louse Infection": Supp Captions

**Supplementary Table 1. Differentially expressed genes of vitamin D-treated salmon.** The tables contain the Ensembl ID, log2 fold change, adjusted p-values, and gene symbols. All of the annotated genes’ accession numbers from BLAST.

**Supplementary Table 2. Quality control data on RNA-seq output.** The table includes information on raw and clean-read counts, base counts, sequencing error rates, quality scores (Q20 and Q30), and GC content for each sample.

**Supplementary Table 3. Novel differentially expressed genes.** Contains all significant (adj p ≤ 0.05) that did not possess an Ensembl ID. The table contains the log2 fold change and adjusted p-values of the novel genes.

**Supplementary Table 4. Differentially expressed immune-related genes of vitamin D-treated salmon.** The tables contain the Ensembl ID, log2 fold change, adjusted p-values, and gene symbols for all identified immune-related genes and annotated via accession numbers from BLAST.

**Supplementary Table 5. Gene set enrichment analysis of vitamin D-treated salmon.** Contains all outputs from GESA with significance of adj p ≤ 0.05 and q value ≤ 0.05. The table contains all ontologies of biological process, molecular function and cellular component.

**Supplementary Table 6. Gene set enrichment analysis of salmon infected with salmon lice.** Contains all outputs from GESA with significance of adj p ≤ 0.05 and q value ≤ 0.05. The table contains all ontologies of biological process, molecular function and cellular component.
